# Improving magnetic resonance spectroscopy in the brainstem periaqueductal grey using spectral registration

**DOI:** 10.1101/2023.03.29.534678

**Authors:** Laura Sirucek, Niklaus Zoelch, Petra Schweinhardt

## Abstract

**Purpose:** Functional understanding of the periaqueductal grey (PAG), a clinically relevant brainstem region, can be advanced using proton magnetic resonance spectroscopy (^1^H-MRS). However, the PAG’s small size and high levels of physiological noise are methodologically challenging. This study aimed to (1) improve ^1^H-MRS quality in the PAG using spectral registration for frequency and phase error correction, (2) investigate whether spectral registration is particularly useful in cases of greater head motion and (3) examine metabolite quantification using literature-based or individual-based water relaxation times.

**Methods:** Spectra were acquired in 33 healthy volunteers (50.1 years, SD=17.19, 18 females) on a 3T Philipps MR system using a point-resolved spectroscopy sequence optimized with very selective saturation pulses (OVERPRESS) and voxel-based flip angle calibration (effective volume of interest size: 8.8×10.2×12.2 mm^3^). Spectra were fitted using LCModel and signal-to-noise ratios (SNR), N-acetylaspartate peak linewidths and Cramér-Rao lower bounds (CRLBs) were measured after spectral registration and after minimal frequency alignment.

**Results:** Spectral registration improved SNR by 5 % (*p*=0.026, median value post-correction: 18.0) and spectral linewidth by 23 % (*p*<0.001, 4.3 Hz), and reduced the metabolites’ CRLBs by 1-15 % (*p*’s<0.026). Correlational analyses revealed smaller SNR improvements with greater head motion (*p*=0.010) recorded using a markerless motion tracking system. Higher metabolite concentrations were detected using individual-based compared to literature-based water relaxation times (*p*’s<0.001).

**Conclusion:** This study demonstrates high-quality ^1^H-MRS acquisition in the PAG using spectral registration. This shows promise for future ^1^H-MRS studies in the PAG and possibly also other clinically relevant brain regions with similar methodological challenges.

## 1. Introduction

The periaqueductal grey (PAG) is a brainstem region with multiple pivotal functions for the human organism including the coordination of cardiovascular, respiratory, motor, and pain modulatory reactions to stress.^1^ Various pathological conditions present with changes in PAG function. For instance, altered PAG functional connectivity has been observed in neurodegenerative diseases,^2,3^ migraine,^4,5^ headache,^6^ fibromyalgia,^7-9^ neuropathic pain,^10^ and chronic low back pain.^11^ A more complete understanding of PAG function in health and disease can be gained by examining the PAG’s neurochemical properties. Proton MR spectroscopy (^1^H-MRS) offers a non-invasive method to obtain in vivo neurochemical information about human brain tissue.

In the PAG, ^1^H-MRS has been investigated in patients with chronic migraine,^12,13^ chronic daily headaches,^14^ and chronic whiplash injury.^15^ Yet, the number of ^1^H-MRS studies in the PAG is limited, which may be due to several difficulties associated with ^1^H-MRS acquisition in this region: first, the PAG is a small structure of approximately 4-5 mm diameter and 14 mm length,^16^ i.e. approximately 5×5×14 mm^3^ (APxLRxFH). Second, the PAG is prone to physiological noise due to its proximity to pulsating anatomical structures, such as major arteries and cerebrospinal fluid (CSF)-filled spaces.^17^ The existing PAG ^1^H-MRS studies employed different approaches to address these challenges. Lai and colleagues^12^ used a long echo time (TE) of 144 ms, which produces a better baseline and allows a more stable detection of the main metabolites creatine (Cre), choline (Cho), and N-acetylaspartate (NAA).^18^ However, various neurobiologically relevant metabolites, e.g. myo-inositol (mI) or glutamate (Glu), require shorter TEs to be detected. Another option is to sacrifice regional specificity by using volume of interest (VOI) sizes lager than the PAG itself, e.g. 20×20×20 mm^3^,^13,14^ because the signal-to-noise ratio (SNR) increases proportionally to VOI size.^19^ Smaller VOIs require longer acquisition times to achieve sufficient SNR, which might in turn impair the spectral quality because of increasing frequency drifts over time and a higher risk for head motion inducing additional frequency and phase errors.^20,21^ Promising tools to correct for frequency and phase drifts or errors are offered by advanced post-processing techniques such as spectral registration.^20^

In this study, the aim was to record high-quality ^1^H-MR spectra in a 8.8×10.2×12.2 mm^3^ VOI covering the PAG of healthy volunteers using a point-resolved spectroscopy sequence (PRESS)^22,23^ optimized with very selective saturation pulses (OVERPRESS)^24-26^ and voxel-based flip angle calibration^27,28^ combined with spectral registration.^20^ The spectral quality was compared to the spectral quality achieved with minimal frequency alignment, i.e. using the unsuppressed water peak of multiple interleaved spectra. In addition, it was investigated whether spectral registration was particularly useful in cases of greater head motion measured with a markerless motion tracking system.^29^ Lastly, because tissue-specific water T_1_ and T_2_ relaxation times vary across different brain regions^30-32^ and standard literature-based values might not generalize to the PAG, differences in metabolite concentrations using literature-based or individual-based water relaxation times were examined. The data presented in this manuscript are part of a larger study investigating differences in PAG spectra between pain-free volunteers and chronic low back pain patients.

## 2. Methods

### 2.1 Participants

Thirty-four healthy volunteers were recruited via online advertisements and oral communication. The participants were age- and sex-matched to a cohort of chronic low back pain patients as part of a larger study (Clinical Research Priority Program “Pain”, https://www.crpp-pain.uzh.ch/en.html). The patient cohort data are not part of the present research question and therefore not discussed. Inclusion criteria were between 18 and 80 years of age and free of low back pain lasting longer than 3 consecutive days during the last year. Exclusion criteria comprised any major medical or psychiatric condition, pregnancy, inability to follow study instructions and any contraindication to MR imaging. The study was approved by the local ethics committee “Kantonale Ethikkommission Zürich” (Nr.: 2019-00136, clinicaltrials.gov: NCT04433299) and was performed according to the guidelines of the Declaration of Helsinki (2013). Written informed consent was obtained from all participants before the start of the experiment.

### 2.2 Study Design Overview

The larger study comprised three experimental sessions of approximately 3 hours each. The first two sessions included clinical, neurophysiological and psychophysical assessments. During the third session, participants underwent two MR measurements, one ^1^H-MRS scan and one resting state functional MR imaging scan with a break of one hour in between. All scans were performed after 12 pm. Only the ^1^H-MRS data are subject of the present study.

### 2.3 Magnetic resonance spectroscopy

Technical details of the ^1^H-MRS acquisition, post-processing and metabolite quantification are listed in Table S1. The following sections provide a brief overview of the applied methods.

#### 2.3.1 Acquisition

^1^H-MRS was acquired on a 3T MR system using a 32-channel receive-only phased-array head coil (Philips Healthcare, Best, The Netherlands). Prior to ^1^H-MRS acquisition, high-resolution (1 mm^3^ isotropic) anatomical T_1_-weighted images were obtained. Based on the 3D T_1_ images, the VOI was placed to cover the PAG according to anatomical landmarks by the same examiner (LS) for all participants (Figure 1). Spectra were localized using a water-suppressed single-voxel PRESS sequence^22,23^ (repetition time (TR): 2500 ms, TE: 33 ms, number of signals averaged: 512 divided into 8 blocks of 64) optimized with six very selective saturation (VSS) pulses (OVERPRESS)^24-26^ to minimize errors in chemical-shift displacement and to achieve consistent localization volumes across all metabolites of interest (Figure 1), and with voxel-based flip angle calibration^27,28^ to achieve an optimal flip angle within the VOI. Accounting for the VSS pulses, the resulting VOI size was 8.8×10.2×12.2 mm^3^=1.1 mL.

**Figure 1.**
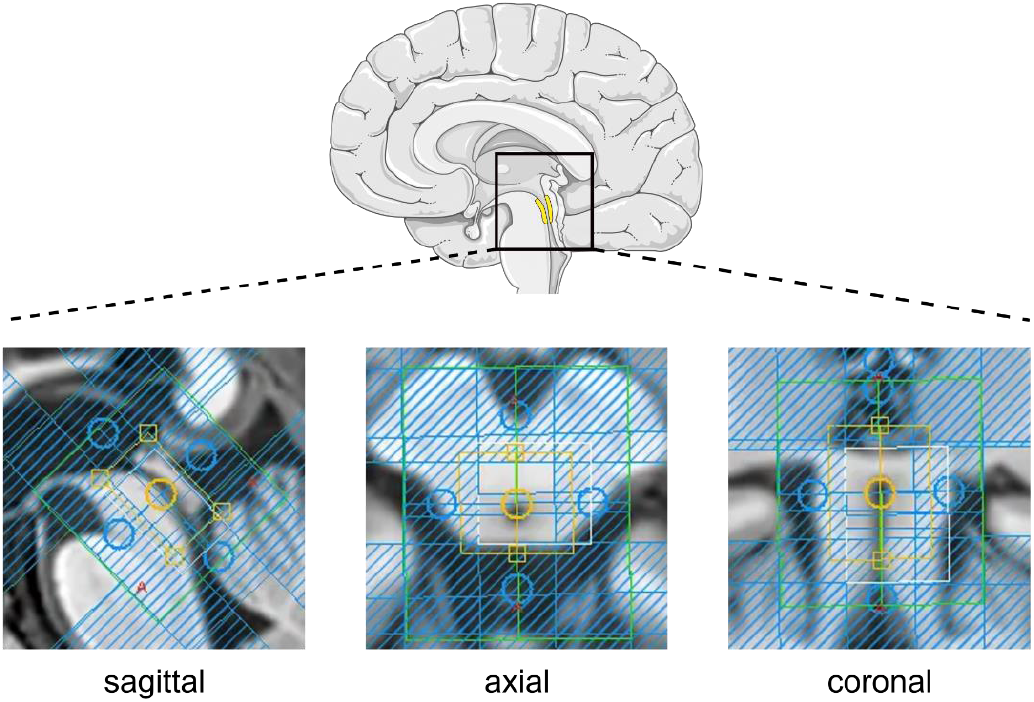
Volume of interest (VOI) placement over the brainstem periaqueductal grey (PAG). The yellow box depicts the VOI’s nominal size (11×15×18 mm^3^ [APxLRxFH]. The blue-shaded areas depict the six very selective saturation (VSS) bands used to minimize errors in chemical-shift displacement and to achieve consistent localization volumes across all metabolites of interest. The unshaded area within the yellow box depicts the final VOI size (8.8×10.2×12.2 mm^3^) accounting for the VSS bands. The schematic brain was adapted from Servier Medical Art (smart.servier.com).

For each individual, two water signals were acquired: one for eddy-current correction and literature-based water referencing obtained from interleaved water unsuppressed spectra (one before each of the 8 blocks) during the ^1^H-MRS acquisition in the PAG (WR-shortTR) and one for an individual-based water referencing approach (see 2.3.3 Metabolite quantification), where a fully-relaxed water signal was estimated from a separate water reference scan after the ^1^H-MRS acquisition in the PAG within the same VOI with a TR of 10000 ms (WR-longTR) and varying TEs (33/66/107/165/261/600 ms). The 3D T_1_ and ^1^H-MRS acquisition in the PAG took 7 min 32 s and 23 min 20 s, respectively.

#### 2.3.2 Post-processing

For the approach *with spectral registration*, frequency alignment was performed using spectral registration in the time domain (adopted from^20,33^). For that, the data was filtered with a 2 Hz Gaussian filter. Only the first 500 ms were used for alignment and the single averages were aligned to the median of all averages. *Without spectral registration*, minimal frequency alignment was achieved by eddy-current correction,^34^ i.e. eddy-current correcting the 64 averages of each block using the unsuppressed water scan (WR-shortTR) acquired prior to the respective block.

For both approaches, the post-processed spectra were visually checked for artefacts. Spectra with artefacts were excluded from further analyses, as well as spectra presenting with insufficient quality,^28^ i.e. with a full width at half maximum (FWHM) value of the unsuppressed water peak (FWHM H_2_O; shim quality indicator) above 2.5 median absolute deviation (MAD)^35^ of the group median or with an SNR (as obtained from LCModel) below 2.5 MAD of the group median.

#### 2.3.3 Metabolite quantification

All spectra were analyzed using LCModel (6.3).^36^ Metabolite concentrations are reported as ratios to the unsuppressed water signal and reflect an estimation of metabolite concentration in moles per kg of tissue water excluding water within CSF. The fully relaxed water signal was estimated in two ways: (1) literature-based, i.e. using the unsuppressed water signal from the WR-shortTR scans and literature values to correct relaxation attenuation. And (2), individual-based, using the TE series acquired with the WR-longTR scan. Varying TEs allow to estimate the T_2_ relaxation time of water within the VOI and therewith, to obtain a subject-specific approximation of the fully-relaxed water signal.^37^ Next to being independent of literature-based tissue-specific T_1_ and T_2_ relaxation times, this approach is also less reliant on correct segmentation of grey matter (GM), white matter (WM) and CSF compared to the literature-based approach.

#### 2.3.4 Motion tracking

Head motion was measured in 25 participants using the markerless motion tracking system Tracoline TCL3 with the TracSuite software 3.1.9 (TracInnovations, Ballerup, Denmark) used for retrospective motion correction of positron emission tomography scans^38,39^ and prospective real-time motion correction of MR imaging scans.^39,40^ A more detailed description of the Tracoline system is provided in Figure S1. A representative recording of a participant’s absolute 3D motion is displayed in Figure 2. From this 3D motion recording, two 3D motion parameters were calculated (Figure 2): (1) the 3D motion standard deviation (3D SD) during the ^1^H-MRS scan, representing an estimate for head motion variability and (2) the difference in 3D motion (3D DIFF) from the beginning to the end of the ^1^H-MRS scan, representing an estimate of the mean head displacement.

**Figure 2.**
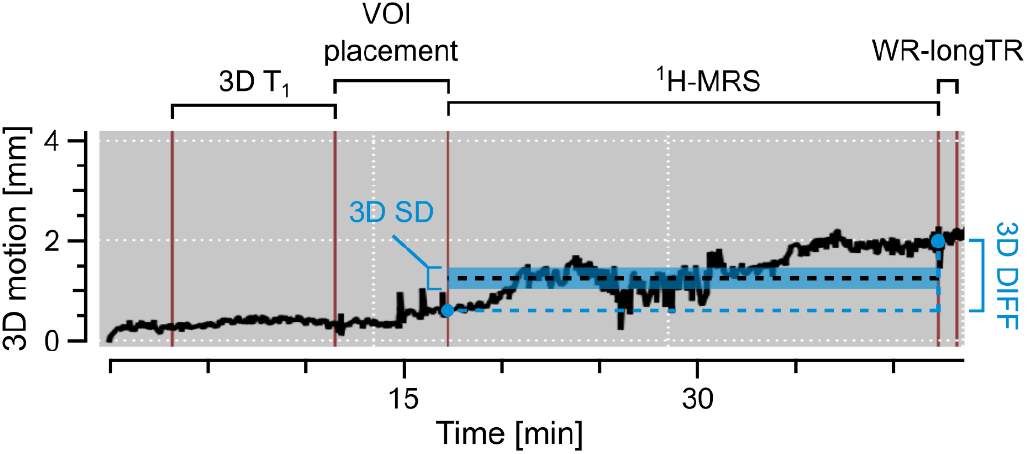
Representative absolute 3D motion recording. The two calculated 3D motion parameters are visually represented in blue: 3D motion standard deviation (3D SD) as estimate for head motion variability and (2) the difference in 3D motion (3D DIFF) as estimate of the mean head displacement. Figure S1 describes how mean head displacement is related to the conventional six-parameter description of head motion, i.e. x-, y-, and z-translations and rotations. The beginning and the end of the proton magnetic resonance spectroscopy (^1^H-MRS) acquisition in the periaqueductal grey (PAG) was manually labelled during the scan (red lines) within the TracSuite software. 3D T_1_: anatomical T_1_-weighted scan; VOI: volume of interest; WR-longTR: water reference scan with a long repetition time.

### 2.4 Statistical analysis

All statistical analyses were performed using RStudio for Mac (2022.12.0+353). Statistical significance was set at α=0.05 with a false discovery rate (FDR) correction per tested research question. The number of corrected tests per research question is indicated as N-FDR. Normal distribution was assessed via inspection of histograms and QQ-plots. Because the majority of investigated outcome measures was not normally distributed, all values are reported as median (interquartile range) and all statistical analyses were performed using non-parametric tests.

SNR and FWHM of the NAA peak (FWHM NAA; as obtained from LCModel; post-processing quality indicator), as well as absolute Cramér-Rao lower bounds (CRLB) (relative CRLBs corrected for the metabolite concentration^41^) for the metabolites tCre (Cre + phosphocreatine), tCho (glycerophospho-choline + phosphocholine), tmI (mI + glycine), tNAA (NAA + N-acetylaspartylglutamate), pooled levels of Glu + Glutamine (Glx), and GABA, were compared between spectra processed with spectral registration and spectra processed without spectral registration using Pratt signed-rank tests (an alternative for Wilcoxon signed-rank tests accounting for ties^42^) (N-FDR=8).

To investigate whether spectral registration was particularly useful in cases of greater head motion, relative improvements in SNR and FWHM NAA were correlated with head motion variability (3D SD) and mean head displacement (3D DIFF) using Spearman correlations (N-FDR=4).

Metabolite concentrations of tCre, tCho, tmI, tNAA, Glx and GABA were compared between the literature-based and the individual-based metabolite quantification approach, i.e. using the water signal from the WR-shortTR scan and using the water signal from the WR-longTR scan, respectively, using Pratt signed-rank tests (N-FDR=6).

For Pratt signed-rank tests, effect sizes are reported as r (small effect: 0.1-<0.3, medium effect: 0.3-<0.5, large effect: ≥0.5 large effect^43^).

## 3. Results

### 3.1 Participant demographics

Out of the 35 recruited participants, one was excluded due to a suspected neurological disorder and one discontinued the scanning session due to discomfort. This resulted in a sample of 33 participants (mean age of 50.1 years, SD=17.19, 18 females) in whom ^1^H-MRS was performed.

For spectra processed with spectral registration and spectra processed without spectral registration, visual inspection led to the exclusion of the same three participants (Figure S2). No participant presented with FWHM H_2_O above or SNR values below 2.5 MAD of the sample median.

### 3.2 Improved spectral quality with spectral registration

A representative single spectrum and overlaid single spectra together with the group average for both post-processing approaches are shown in Figure 3. The FWHM H_2_O reflecting the quality of the shim was 5.4 Hz (4.88-5.62) (Figure 4A).

**Figure 3.**
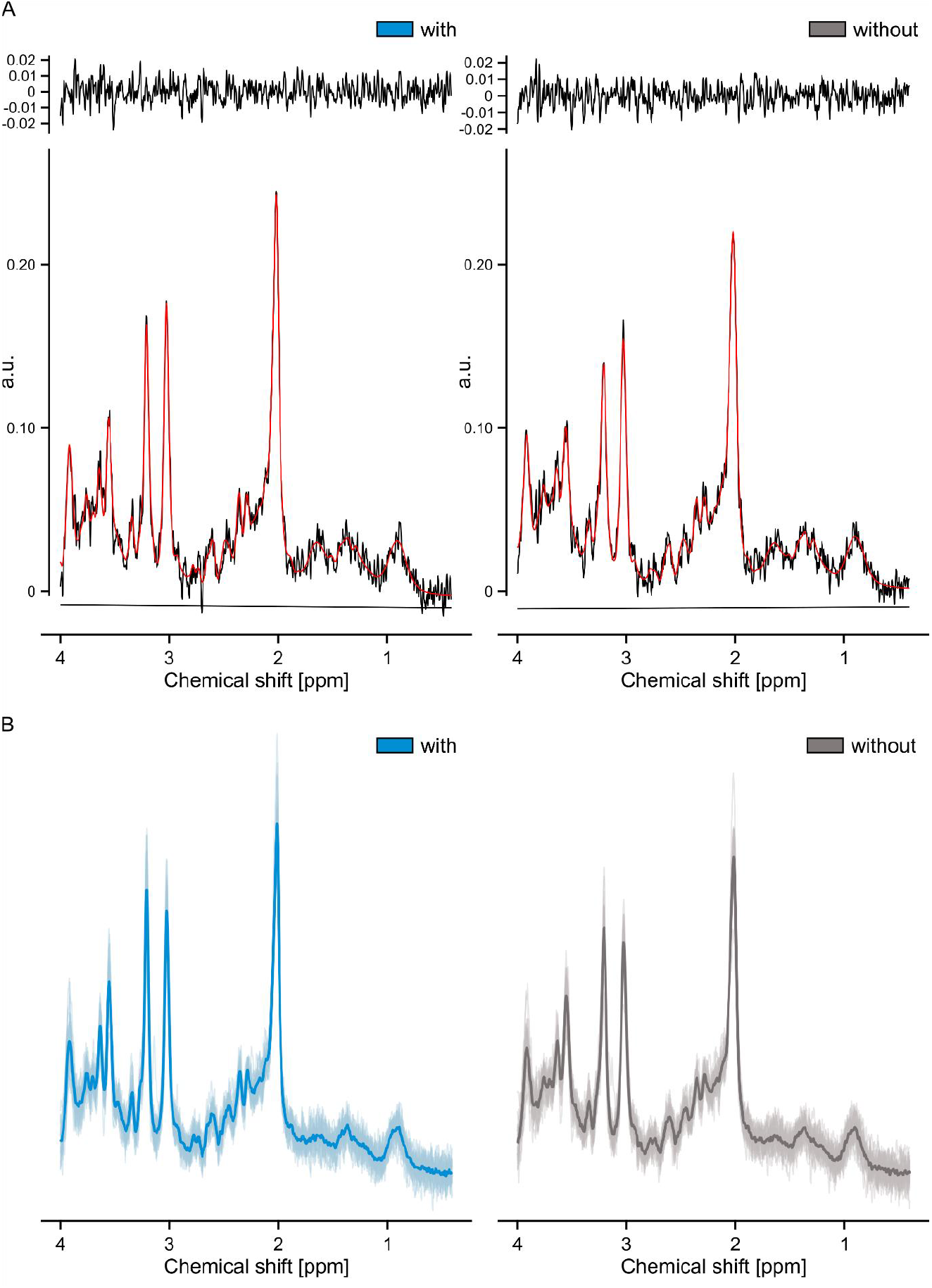
Spectra processed with and without spectral registration. Representative single spectra and overlaid LCModel fits (red line) shown in (A) are from the same individual who showed a signal-to-noise ratio of 18, i.e. the group median, in both post-processing approaches and an N-acetylaspartate linewidth improvement from 6.9 Hz without spectral registration (right panel) to 4.9 Hz with spectral registration (left panel) (29.6%). (B) Overlaid single spectra together with the group average (bold) for spectra processed with (blue) and spectra processed without (grey) spectral registration. Spectra were acquired using a point-resolved spectroscopy sequence optimized with very selective saturation pulses (OVERPRESS) and voxel-based flip angle calibration (TR: 2500 ms, TE: 33 ms, number of signals averaged: 512 divided into 8 blocks of 64, volume of interest size: 8.8×10.2×12.2 mm^3^=1.1 mL).

**Figure 4.**
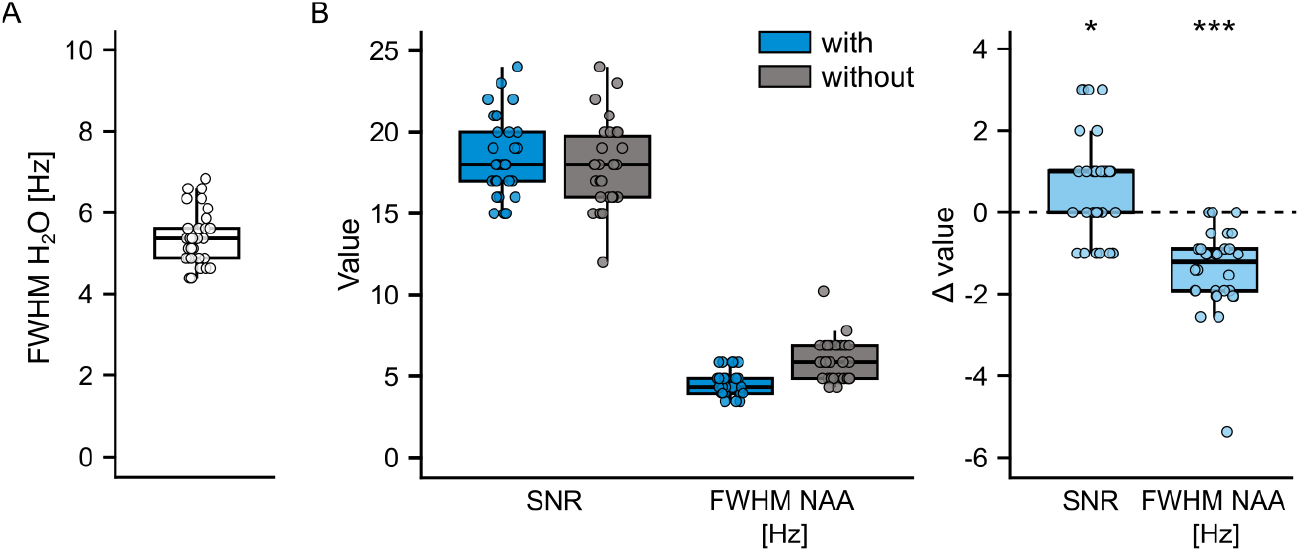
Improved spectral quality with spectral registration. Full width at half maximum values of the water peak (FWHM H_2_O) (A) reflect the quality of the shim and apply to both post-processing approaches. Differences (Δ) in SNR and full width at half maximum values of the N-acetylaspartate peak (FWHM NAA) were calculated by subtracting the values obtained without spectral registration from the values obtained with spectral registration. An improvement using spectral registration is reflected in positive differences for SNR and in negative differences for FWHM NAA. * *p*<0.05, *** *p*<0.001.

Spectral registration significantly improved the SNR (median improvement: 4.8 %, Z=2.26, *p*=0.026, r=0.41), FWHM NAA (22.6 %, Z=-4.72, *p*<0.001, r=0.86), and absolute CRLBs of all investigated metabolites (1.3-14.7 %, Z’s<-2.22, *p*’s<0.026, r’s>0.41) of the acquired spectra (Table 1, Figure 4B). Given the low SNR of the single averages, spectral registration required a sufficiently large residual water peak (ratio to NAA peak: 8.8 [8.40 - 9.60]).

**Table 1.** Quality measures and metabolite concentrations for ^1^H-MRS spectra acquired in the PAG.

| Measure | with spectral registration | without spectral registration | difference | difference [%] | Z | p | effect size r |
| --- | --- | --- | --- | --- | --- | --- | --- |
| SNR | 18.0 (17.00-20.00) | 18.0 (16.00-19.75) | 1.0 (0.00-1.00) | 4.8 (0.00-6.25) | 2.26 | 0.026 | 0.41 |
| FWHM NAA [Hz] | 4.3 (3.96-4.85) | 5.9 (4.85-6.90) | -1.2 (-1.92--0.90) | -22.6 (-29.63--14.81) | -4.72 | <0.001 | 0.86 |
| CRLB tCre [I.U.] | 22.6 (21.78-24.50) | 23.0 (22.08-30.80) | -0.3 (-0.67-0.12) | -1.3 (-3.10-0.57) | -2.22 | 0.026 | 0.41 |
| CRLB tCho [I.U.] | 9.9 (7.72-10.93) | 10.7 (9.38-11.85) | -0.2 (-0.92--0.09) | -2.5 (-7.5--0.89) | -3.96 | <0.001 | 0.72 |
| CRLB tml [I.U.] | 38.0 (31.33-41.37) | 39.9 (34.72-43.05) | -0.4 (-1.03--0.14) | -1.3 (-2.82--0.43) | -2.49 | 0.017 | 0.45 |
| CRLB tNAA [I.U.] | 29.5 (28.42-30.60) | 30.0 (28.54-32.11) | -0.4 (-0.62-0.08) | -1.3 (-2.10-0.24) | -2.72 | 0.013 | 0.50 |
| CRLB Glx [I.U.] | 83.6 (77.84-89.25) | 85.5 (81.34-94.10) | -4.1 (-6.62-0.32) | -4.7 (-6.83-0.37) | -2.53 | 0.017 | 0.46 |
| CRLB GABA [I.U.] | 62.44 (52.29-66.35) | 72.47 (64.11-77.51) | -10.4 (-14.00--3.93) | -14.7 (-19.18--6.50) | -4.77 | <0.001 | 0.87 |
| CRLB tCre [%] | 2 (2-2) | 2 (2-3) |  |  |  |  |  |
| CRLB tCho [%] | 3 (2-3) | 3 (3-3) |  |  |  |  |  |
| CRLB tml [%] | 3 (2-3) | 3 (3-3) |  |  |  |  |  |
| CRLB tNAA [%] | 2 (2-2) | 2 (2-2) |  |  |  |  |  |
| CRLB Glx [%] | 5 (5-6) | 5 (5-6) |  |  |  |  |  |
| CRLB GABA [%] | 24 (20-27) | 21.5 (17.25-28.75) |  |  |  |  |  |
|  | WR-shortTR water signal | WR-longTR water signal | difference | difference [%] | Z | p | effect size r |
| tCre [mmol/kg] | 6.2 (6.05-6.45) | 6.7 (6.42-6.93) | 0.5 (0.36-0.61) | 7.5 (5.75-9.21)<sup>†</sup> | 4.35 | <0.001 | 0.79 |
| tCho [mmol/kg] | 2.1 (1.93-2.18) | 2.2 (2.08-2.35) | 0.2 (0.12-0.20) |  | 4.35 | <0.001 | 0.79 |
| tml [mmol/kg] | 7.7 (7.14-8.07) | 8.2 (7.74-8.64) | 0.6 (0.45-0.74) |  | 4.31 | <0.001 | 0.79 |
| tNAA [mmol/kg] | 8.3 (8.15-8.56) | 8.9 (8.56-9.44) | 0.6 (0.47-0.76) |  | 4.39 | <0.001 | 0.80 |
| Glx [mmol/kg] | 9.2 (8.49-10.13) | 9.7 (9.06-10.73) | 0.7 (0.52-0.89) |  | 4.29 | <0.001 | 0.78 |
| GABA [mmol/kg] | 1.5 (1.26-1.67) | 1.6 (1.33-1.83) | 0.1 (0.06-0.15) |  | 4.41 | <0.001 | 0.81 |

#### Less improvement in spectral quality with greater head motion

On average, participants showed a head motion variability (3D SD) of 0.5 mm (0.32-0.73) and a mean head displacement (3D DIFF) of 1.7 mm (0.62-2.64) during the ^1^H-MRS scan.

Greater mean head displacement (3D DIFF; rho=-0.58, *p*=0.010) but not head motion variability (3D SD; rho=-0.31, *p*=0.259) was associated with less relative improvement in SNR (Figure S3). Relative improvement in FWHM NAA was not associated with head motion measures (rho’s <0.3, *p*’s >0.259).

### 3.3 Higher metabolite concentrations using individual-based quantification approach

Metabolite quantification using the subject-specific water signal from the WR-longTR scan, i.e. the individual-based quantification approach, yielded higher concentrations for all investigated metabolites (Z’s >4.29, *p*’s<0.001, r’s>0.78) compared to the literature-based approach using the water signal from the WR-longTR scan (Table 1, Figure S4). T_2_ relaxation times estimated via the TE series in the WR-longTR scan were 81.25 ms (77.75-84.2).

## 4. Discussion and Conclusions

This study aimed to record high-quality ^1^H-MR spectra with optimized regional specificity in the brainstem PAG. Spectral registration^20^ was used to address the challenge of increased frequency drifts and phase errors during the prolonged ^1^H-MRS acquisition needed for the small VOI size of 8.8×10.2×12.2 mm^3^. Using spectral registration, all quality measures of the acquired spectra improved, resulting in high spectral quality in the methodologically challenging PAG.

Compared to previous PAG ^1^H-MRS studies,^12-15^ this study’s approach allowed higher regional specificity together with quantification of neurobiologically relevant metabolites detected at short TEs such as Glx. Because these previous studies did not report spectral quality measures, a direct quality comparison is not possible. However, the here achieved spectral quality is comparable or even superior to studies which examined other brainstem VOIs of larger size^44-49^ and superior to one study with a brainstem VOI of similar size.^50^ Spectral registration improving SNR and NAA linewidth by approximately 5% and 23%, respectively, is relevant for several reasons. First, using spectral registration, VOI size could be decreased by 5% or acquisition time by 10%. This moderate effect is complemented by the more pronounced linewidth improvement. Narrower linewidths benefit metabolite detection independent of SNR, e.g. it has been shown that at constant SNR, broader linewidths decrease metabolite quantification accuracy.^51^ This is most likely due to hampered peak fitting and differentiation of overlapping peaks, supported by observations of larger absolute CRLBs with broader linewidths.^52^ Glutamate quantification appears to be highly susceptible to linewidth broadening^53^ and thus, narrow linewidths might particularly benefit ^1^H-MRS focusing on excitatory neurotransmitter levels. The here reported linewidth improvement is approximately 10% lower compared to the original report of spectral registration effects which showed a linewidth reduction of the residual water peak by 31.7%.^20^ This is probably explained by the fact that in the present study, spectral registration was compared to minimal frequency alignment using the unsuppressed water peak of multiple interleaved spectra, while in the original report, linewidths were compared before and after spectral registration. Regarding GABA, PRESS sequences have limitations because GABA’s signal overlaps with resonances from other metabolites and is therefore difficult to detect. Other techniques allow a more reliable GABA detection, such as the MEscher-GArwood (MEGA) PRESS.^54^ Yet, MEGA-PRESS is highly susceptible to frequency drifts^55^ and requires VOI sizes of approximately 30×30×30 mm^3^, making it unsuitable for the PAG. Moreover, the CRLBs for GABA of the present study (median absolute: 62.44 I.U., median relative: 24 %) indicate an adequate reliability of the concentration measurement and the detected concentrations lie within the expected range for human brain tissue.^56^ A disadvantage of the sequence used here is the prolonged scan duration, which might be particularly challenging for patient cohorts. Advances in MR methodology, e.g. higher field strengths, might allow to achieve similar spectral quality with shorter scan durations. Nevertheless, the presented technique offers new opportunities to investigate PAG function and might translate to other clinically relevant small-sized brain regions prone to physiological noise, such as the hippocampus.

Head motion is a source for frequency and phase errors.^20,21^ Because spectral registration corrects for frequency and phase errors,^20^ it might be particularly useful in cases of greater head motion. Interestingly, the opposite was observed, i.e. the more participants moved during the ^1^H-MRS scan, the smaller the SNR improvement. Potential reasons for this observation were examined by exploring how spectral quality improvement and head motion were associated with measured frequency drifts (displayed in Figure S5) and phase errors. It was observed that (1) spectral registration mainly improved SNR for spectra with larger frequency fluctuations but not phase fluctuations, and (2) larger phase fluctuations were associated with smaller FWHM NAA improvement (Figure S6). Thus, in the present study, spectral registration corrected for frequency errors while phase errors impeded effective spectral registration. Head motion did not correlate with frequency drifts or phase errors, but previous studies have shown that translational head motion leads to phase shifts while frequency changes in response to small head movements may be negligible.^21^ Taken together, head motion might have induced phase errors which spectral registration was not able to account for, resulting in negative effects on spectral quality improvement. Of note, the here used motion tracking system is based on 3D surface tracking, which cannot be directly translated to the actual displacement of the PAG. Thus, it is possible that smaller scale motion, e.g. physiological motion of the brainstem, was corrected for by spectral registration. Overall, the markerless motion tracking system allows simple real-time head motion monitoring which might help decisions on whether a sequence should be repeated due to excessive head motion.

The current gold-standard for metabolite quantification is referencing the metabolite signal to the water signal from the same VOI corrected for partial volume and tissue-specific relaxation effects.^57^ Literature provides reference values for the therefore required tissue-specific water T_1_/T_2_ relaxation times, but these might not generalize to the PAG because T_1_/T_2_ relaxation times vary across different brain regions.^30-32^ Therefore, metabolite concentrations were not only quantified using literature-based (WR-shortTR) but also individual-based (WR-longTR) water relaxation times. Using the individual-based approach, 7.5% higher metabolite concentrations were estimated compared to the literature-based approach meaning that the estimated water signal was smaller when individual-based water relaxation times were used. This effect is most likely due to T_1_ relaxation time differences between the two approaches, because the here used literature-based T_2_ relaxation times were similar to the T_2_ values estimated via the TE series in the WR-longTR scan. This result raises the question whether the standard T_1_ relaxation times apply to the PAG. Regardless, the “absolute” metabolite concentrations reported are valuable in that they allow comparison to future PAG ^1^H-MRS studies.

In summary, spectral registration enabled the acquisition of high-quality ^1^H-MR spectra in the PAG, a physiologically and clinically highly relevant brainstem region. This approach offers the opportunity to further investigate the PAG’s neurochemical properties in health and disease and might not only be applicable in the PAG, but also in other brain regions with similar methodological challenges.

## Data availability

Data will be made available upon request for participants who gave informed consent on further use of their anonymized data.

## Acknowledgments

This study was funded by the Clinical Research Priority Program “Pain” of the University of Zurich. L. Sirucek is supported by the Theodor und Ida Herzog-Egli Stiftung. The authors express their gratitude to Emma Louise Kessler, MD for her generous donation to the Zurich Institute of Forensic Medicine, University of Zurich, Switzerland. We thank all participants who took part in the study. Additionally, we thank Lucas Tauschek, Simon Carisch, Madeleine Hau and Alexandros Guekos for their support during data acquisition.

