## Supporting information for "Improving magnetic resonance spectroscopy in the brainstem periaqueductal grey using spectral registration"

**Table S1.** Technical details of the proton magnetic resonance spectroscopy ($^{1}$H-MRS) acquisition using the experts' consensus checklist for a single voxel $^{1}$H-MRS study[1].

| | |
|---|---|
| **1. Hardware** | |
| a. Field strength [T] | 3 T |
| b. Manufacturer | Philips |
| c. Model (software version if available) | Achieva with dStream Upgrade. Software Release 5.6.1. |
| d. RF coils: nuclei (transmit/receive), number of channels, type, body part | 32-channel dStream receive-only phased-array head coil |
| e. Additional hardware | N/A |
| **2. Acquisition** | |
| a. Pulse sequence | PRESS in combination with six saturation pulses (OVERPRESS)[2-4] and a voxel-based flip angle calibration performed prior to each measure-ment to achieve the desired flip angle and thus optimal SNR.[5,6] |
| b. Volume of Interest (VOI) locations | Periaqueductal grey |
| c. Nominal VOI size [cm$^3$, mm$^3$] | 11x15x18 mm$^3$ (APxRLxFH). Accounting for the saturation pulses, the resulting effective VOI size was 8.8x10.2x12.2 mm$^3$=1.1 mL. |
| d. Repetition Time (TR), Echo Time (TE) [ms, s] | TR: 2500ms, TE: 33ms |
| e. Total number of Excitations or acquisitions per spectrum | 512 acquisitions per spectrum divided into 8 blocks of 64 averages each. At the beginning of each block, one scan without water suppression was performed, resulting in a total of 8 acquisitions for the water reference (WR-shortTR).<br><br>Further, after completion of the 8 blocks, an additional water reference scan with TR 10000ms was performed within the same VOI (WR-longTR). TEs were varied for the 6 acquired averages (+ 2 dummy scans), i.e. 33/66/107/165/261/600 ms, allowing to estimate the $T_2$ relaxation time of water within the VOI and therewith, obtain a subject-specific approxi-mation of the fully-relaxed water signal within the VOI.[7] All settings were kept identical to the previous sequence except the center frequency of the applied pulses which was set to the resonance frequency of water instead of creatine. |

| | |
|---|---|
| f. Additional sequence parameters (spectral width in Hz, number of spectral points, frequency offsets) | Spectral width 2000 Hz, 2048 points. |
| g. Water Suppression Method | VAriable Power radiofrequency pulses with Optimized Relaxation delays (VAPOR) |
| h. Shimming Method, reference peak, and thresholds for “acceptance of shim” chosen | Second-order automatic pencil-beam shimming where pencil-beam excitations were performed trough a shim volume of 30x30x30 $mm^3$ aligned with the spectroscopy VOI (PB volume option). |
| i. Triggering or motion correction method<br><br>(respiratory, peripheral, cardiac triggering, incl. device used and delays) | N/A |
| 3. Data analysis methods and outputs | |
| a. Analysis software | ReconFrame (GyroTools LLC, Zurich, Switzerland) to pre-process the spectra and LCModel version 6.3[8] for analysis. For the pre-processing, code from FID-A[9] was added to ReconFrame. |
| b. Processing steps deviating from quoted reference or product | Processing of .raw/.lab with ReconFrame including a) eddy current correction[10] and b) coil combination.<br><br>Then, for the processing with spectral registration:<br>Frequency alignment using spectral registration in the time domain[11] (adopted from FID-A[9]). For the spectral registration in the time domain, data was filtered with a 2 Hz Gaussian filter. Only the first 500 ms were used for alignment and the single averages were aligned to the median of all averages.<br><br>Without spectral registration:<br>Minimal frequency alignment was achieved by the performed eddy current correction with the interleaved water unsuppressed scans (WR-shortTR scan).<br><br>Both approaches were followed by the following steps:<br>c) residual water filtering, d) 1Hz Gaussian filtering, and e) measurement of FWHM of water peak (FWHM $H_2O$) (Method 1 in FID-A[9]) using 8-fold zero- filling and taking the absolute value of the time domain water signal before fast Fourier transformation. Only the signal from the second unsuppressed water peak from WR-shortTR) was used to determine FHWM $H_2O$. |

| | |
|---|---|
| c. Output measure<br><br>(e.g. absolute concentration, institutional units, ratio) | Ratio to water signal from WR-shortTR or WR-longTR. Ratios were corrected for CSF fraction and multiplied with the inverse of the molecular weight of water. Relaxation attenuation of the metabolite signals was not corrected. With that, a rough estimate of moles of metabolite per mass of tissue water (excluding CSF) - molar concentration mol/kg, was achieved.<br><br>Based on the different WR scans, the fully relaxed water signal was estimated differently:<br><br>WR-shortTR:<br>The WR-shortTR water scan was provided as water reference to LCModel. Relaxation attenuation of the water signal was considered based on literature values. The following $T_1$ /$T_2$ values were used for the different tissue types (ms): GM: 1820 / 100; WM: 1080 / 70; CSF: 4160 / 500 and the following relative densities of NMR-visible water: GM: 0.78, WM: 0.65, CSF: 0.97.<br><br>WR-longTR:<br>In this case the WR-longTR water scan was provided as water reference to LCModel. The fully-relaxed water signal was estimated based on the measured subject-specific TE series. The decay of the water was fitted with an exponential decay within MATLAB 2022 using “fitnlm” and used to estimate the water peak area at TE=0 ms. The ratio of the water peak area at TE=33 ms and TE=0 ms was used to correct the conc. values resulting from LCModel.<br><br>General:<br><br>Relative tissue type volume fractions within the VOI were determined using the $T_1$-weighted planning images (three-dimensional magnetization-prepared rapid gradient-echo (MPRAGE) sequence,[12] 1mm$^3$ isotropic, TE=3.7 ms, TR=8.1 ms, TI=1024 ms, shot interval=3000 ms, FOV: 240x160x240 mm$^3$ (APxLRxFH), flip angle=8°, scan time=7 min32 s) which were segmented using SPM12.[13]<br><br>For both approaches, based on WR-shortTR and based on WR-longTR, ratios to water signal were obtained from LCModel with WCONC=55556 and ATTH2O=1. |
| d. Quantification references and assumptions, fitting model assumptions | The unsuppressed water peak (WR-shortTR or WR-longTR) was used as reference. |

| | |
|---|---|
| | A simulated basis set (simulated using the GAMMA Simulation Package[14]) containing the following 20 metabolites was used to determine peak areas in the chemical shift range from 0.4 ppm and 4.0 ppm:<br>alanine, aspartate, glucose, creatine, phosphor-creatine, glutamine, glutamate, glycerol-phosphocholine, phosphocholine, lactate, myo-inositol, N-acetylaspartate, N-acetylaspartyl-glutamate, scyllo-inositol, glutathione, taurine, glycine, phosphoethanolamine, ascorbate, and γ-aminobutyric acid.<br><br>Simulated contribution of macromolecules and lipid signals were provided within LCModel. |
| 4. Data Quality | |
| a. Reported variables<br><br>(SNR, Linewidth (with reference peaks)) | SNR and FWHM of the N-acetylaspartate peak obtained from the LCModel output. To assess the shim quality in the VOI, FWHM $H_2O$ from the water reference scan (WR-shortTR) was determined. |
| b. Data exclusion criteria | Visual inspection of artifacts and spectra with FWHM $H_2O$ values above 2.5 mean absolute deviance (MAD)[15] of the group median or SNR values below 2.5 MAD of the group median. |
| c. Quality measures of postprocessing Model fitting (e.g. CRLB, goodness of fit, SD of residual) | Absolute CRLBs[16] of selected metabolites, i.e. relative % CRLBs obtained from LCModel multiplied by the conc. values obtained from LCModel. |
| d. Sample Spectrum | Figure 3 |

CRLB: Cramér-Rao lower bound; CSF: cerebrospinal fluid; FOV: field of view; FWHM: full width at half maximum; GM: grey matter; NMR: nuclear magnetic resonance; PRESS: point-resolved spectroscopy; SNR: signal-to-noise ratio; TE: echo time; TR: repetition time; VOI: volume of interest; WM: white matter.

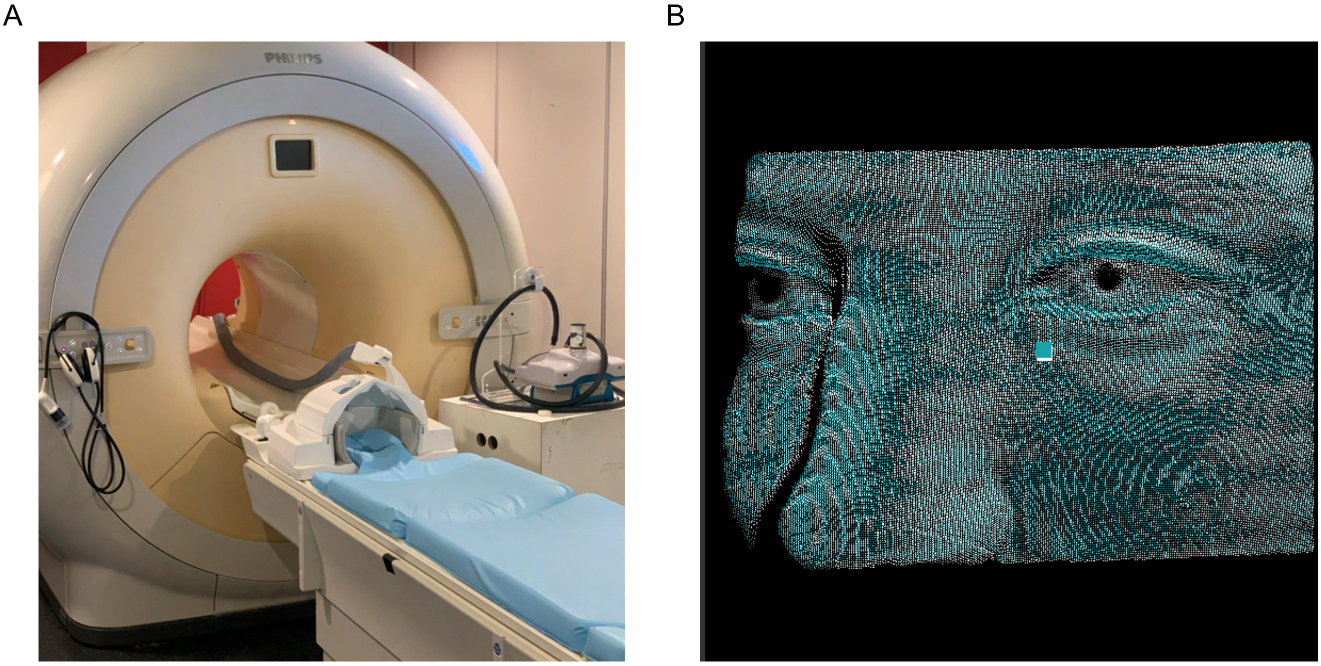

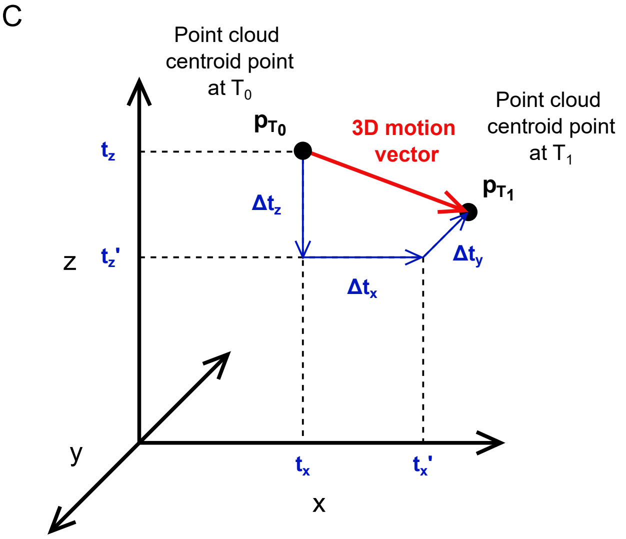


**3D motion vector** $= \left\|p_{T_1} - p_{T_2}\right\|_2 = \left\|p_{T_1} * R_{T_1} + T_{T_2} - p_{T_2}\right\|_2$

With:

$$T = [t_x, t_y, t_z]'$$

$$R = R_x * R_y * R_z$$

$$R_x(\theta) = \begin{bmatrix} 1 & 0 & 0 \\ 0 & \cos\theta & -\sin\theta \\ 0 & \sin\theta & \cos\theta \end{bmatrix}$$

$$R_y(\theta) = \begin{bmatrix} \cos\theta & -\sin\theta & 0 \\ \sin\theta & \cos\theta & 0 \\ 0 & 0 & 1 \end{bmatrix}$$

$$R_z(\theta) = \begin{bmatrix} \cos\theta & -\sin\theta & 0 \\ \sin\theta & \cos\theta & 0 \\ 0 & 0 & 1 \end{bmatrix}$$

**Figure S1**. Details on the markeless motion tracking system Tracoline TCL3 (TracInnovations, Ballerup, Denmark). The Tracoline is a 3D stereo vision system which uses structured invisible infrared light to construct 3D point clouds of the participant's face. (A) The set-up of the Tracoline system. (B) A 3D point cloud reconstruction example (for participant anonymity purposes of author LS's face). The TracSuite software estimates head motion by setting a reference point cloud to which subsequent point clouds are registered to. The calculated 3D motion represents the absolute motion of the point cloud's centroid point. (C) Schematic representation (left panel) and mathematical description (right panel) of how 3D motion relates to the conventional six-parameter description of head motion, i.e. x-, y-, and z- translations and rotations. 3D motion reflects the length of the translation vector between the point cloud centroid point at its starting position $T_0$ and the point cloud centroid point at $T_1$. The schematic contains only translations because rotations cannot be accurately illustrated in this type of schematic.

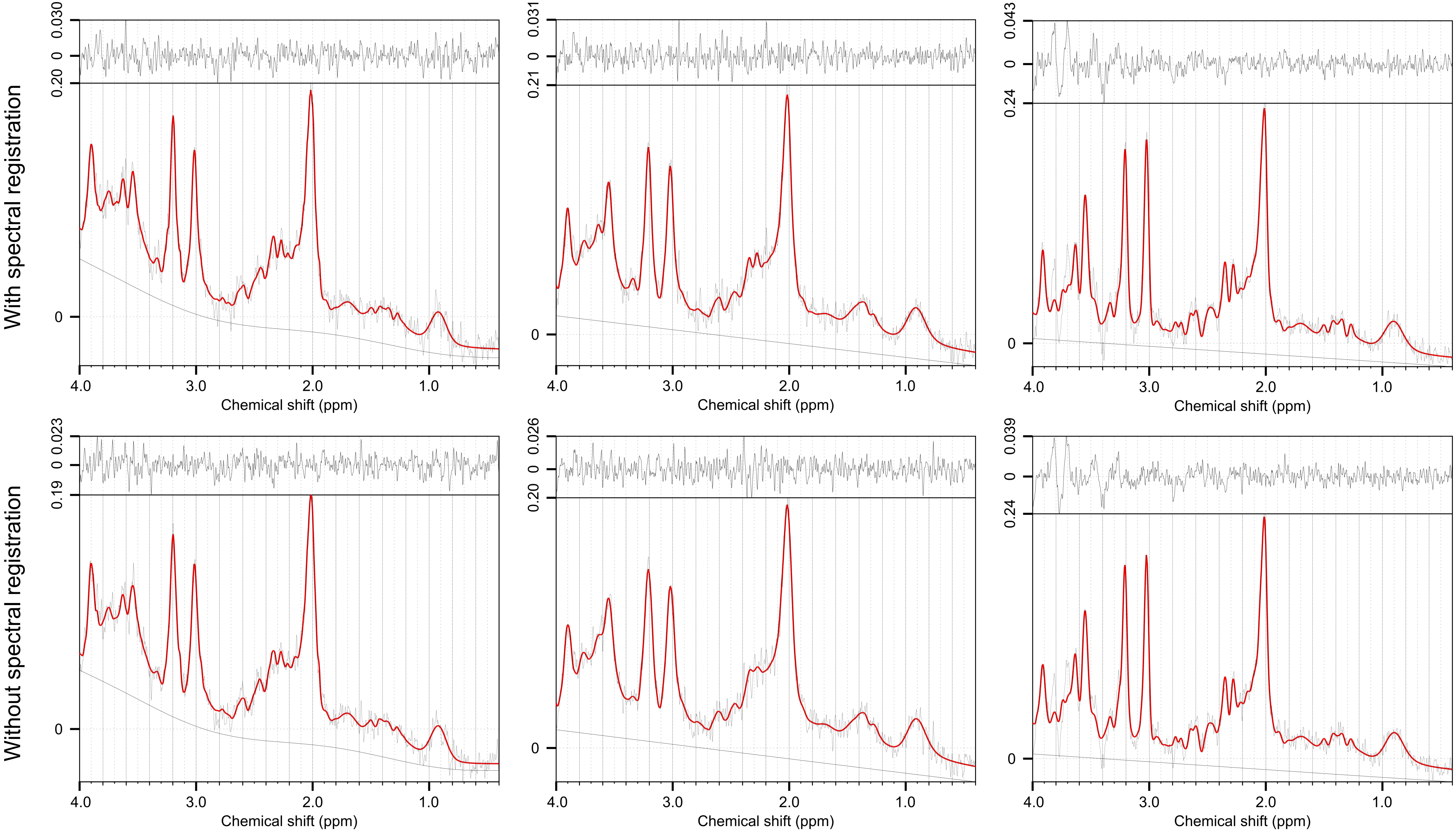


**Figure S2**. Excluded proton magnetic resonance spectra acquired in the brainstem periaqueductal grey. Based on visual inspection for artefacts, the same three spectra were excluded for spectra processed with spectral registration and spectra processed without spectral registration, i.e. minimal frequency alignment using interleaved unsuppressed water peaks.

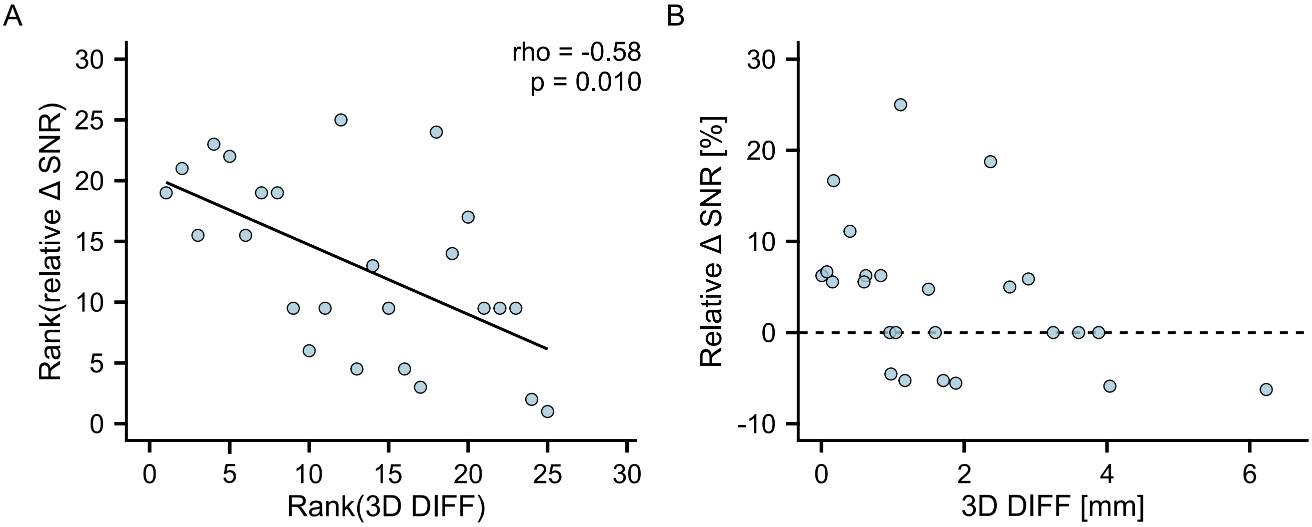


**Figure S3**. Less improvement in spectral quality with greater head motion. Because the majority of investigated outcome measures was not normally distributed, a Spearman's correlation was performed on the ranks of the relative signal-to-noise ratio (SNR) differences (Δ; SNR with spectral registration – SNR without spectral registration divided by SNR without spectral registration; positive values reflecting an SNR improvement using spectral registration) and mean head displacement (3D DIFF) (A). To help with interpretability, the raw data is shown in (B).

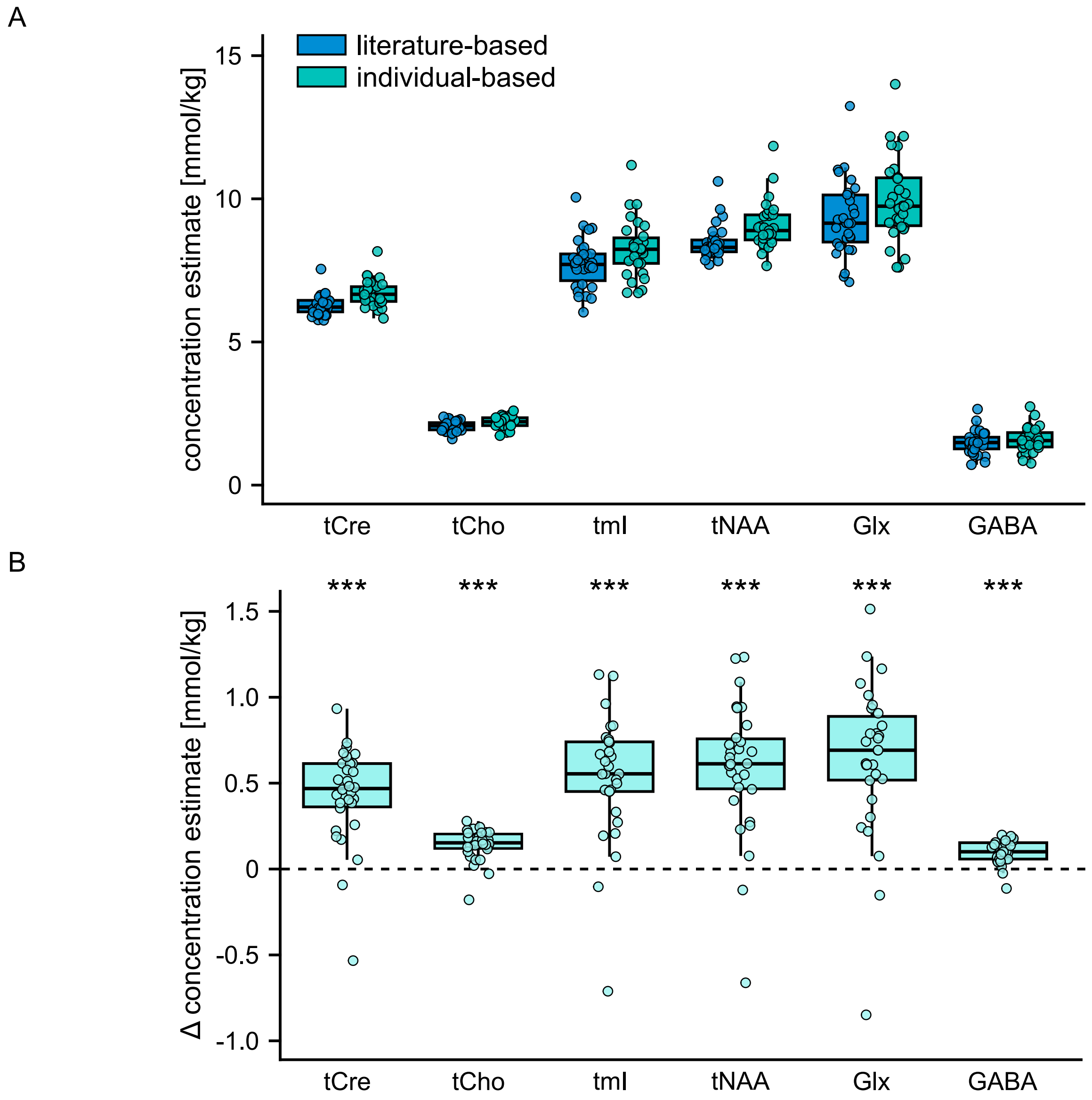


**Figure S4.** Higher metabolite concentrations using individual-based water relaxation times. Differences (Δ) in metabolite concentration estimates were calculated by subtracting the values obtained with literature-based water relaxation times from the values obtained with individual-based water relaxation times. Positive differences reflect higher metabolite concentrations individual-based water relaxation times. *** $p<0.001$. GABA: γ-aminobutyric acid; Glx: glutamate + glutamine; tCho: glycerophosphocholine + phosphocholine; tCre: creatine + phosphocreatine; tmI: myo-inositol + glycine; tNAA: N-acetylaspartate + N-acetylaspartylglutamate.

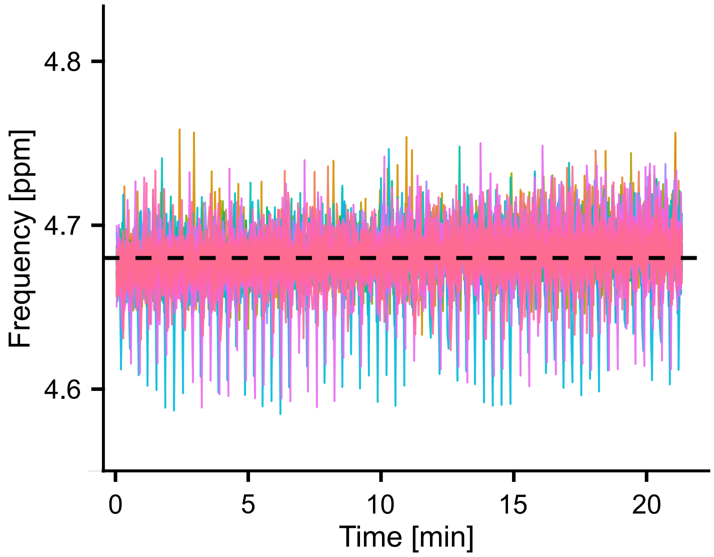


**Figure S5**. Frequency drift as determined with spectral registration shown for all measured subjects over the whole $^{1}$H-MRS acquisition. To allow comparison with previous reports of frequency drifts on Philips MR systems (e.g. [58]) the values obtained in Hz were converted to ppm (- value in [Hz] / 127 + 4.68). The observed overall frequency drifts are comparable to previously published data,[58] except for larger subject-specific variations which were most likely due to the smaller VOI size used in the present study. The dashed line represents the nominal water frequency (4.68 ppm).

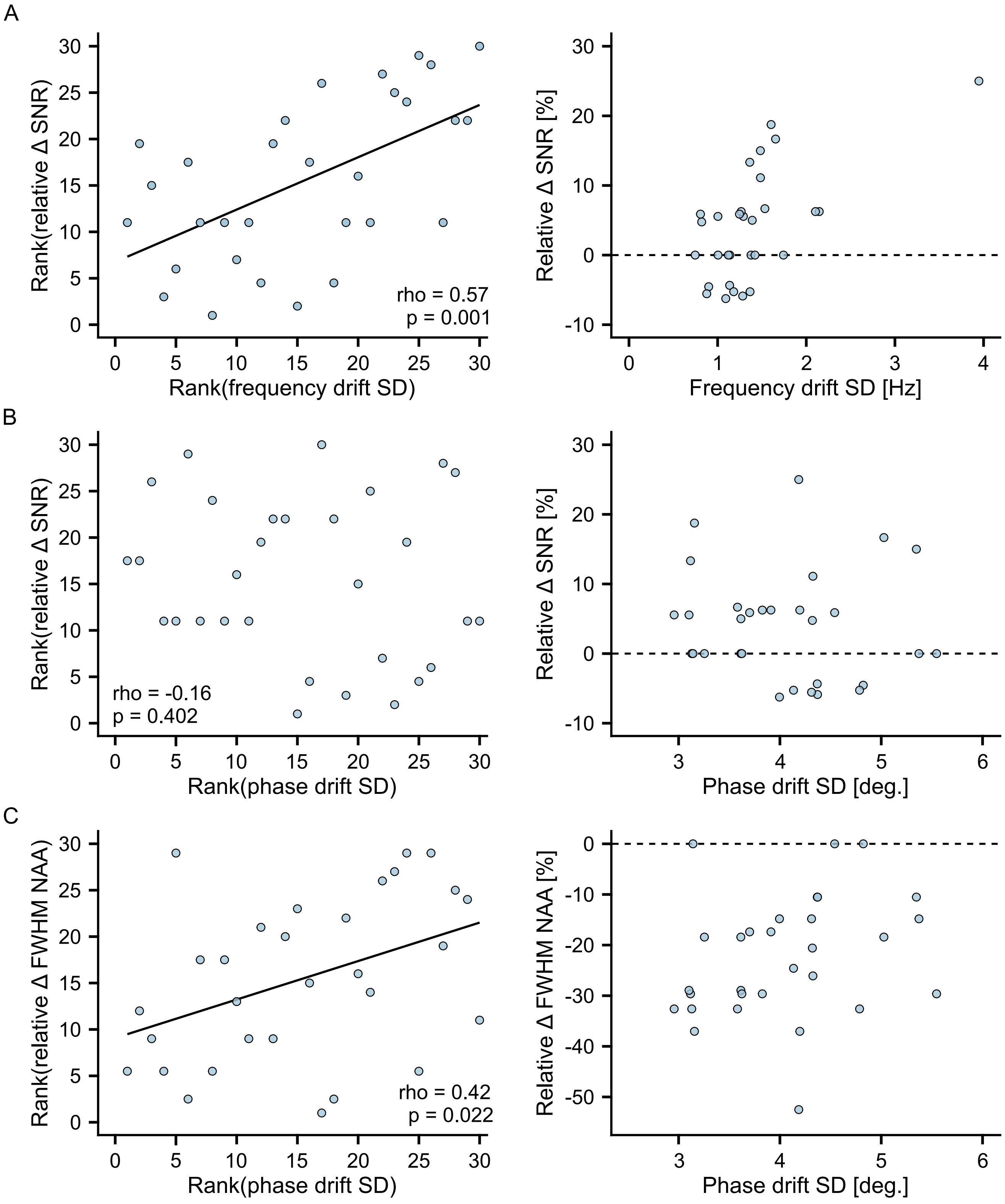


**Figure S6.** Associations of spectral quality improvement with frequency drifts and phase errors. Standard deviations (SD) of measured frequencies and phases served as measure of frequency and phase fluctuations, respectively. (A) Association of signal-to-noise ratio (SNR) improvement with frequency fluctuations. (B) No association of SNR improvement with phase fluctuations. (C) Association of full width at half maximum of the N-acetylaspartate peak (FWHM NAA) improvement with phase fluctuations. Relative differences (Δ): value with spectral registration – value without spectral registration divided by value without spectral registration. An improvement using spectral registration is reflected in positive Δ for SNR and in negative Δ for FWHM NAA. Because the majority of investigated outcome measures was not normally distributed, Spearman's correlations were performed. Left panels show the ranked data and, to help with interpretability, the raw data is shown in the right panels. Multiple comparison correction was not performed because of the exploratory nature of this analysis.
